# Healthier decisions in the cued-attribute food choice paradigm have high test-retest reliability across five task repetitions over 14 days

**DOI:** 10.1101/2021.03.25.437001

**Authors:** Zahra Barakchian, Anjali Raja Beharelle, Todd A. Hare

## Abstract

Food choice paradigms are commonly used to study decision mechanisms, individual differences, and intervention efficacy. Here, we measured behavior from twenty-three healthy young adults who completed five repetitions of a cued-attribute food choice paradigm over two weeks. This task includes cues prompting participants to explicitly consider the healthiness of the food items before making a selection, or to choose naturally based on whatever freely comes to mind. We found that the average patterns of food choices following both cue types and ratings about the palatability (i.e. taste) and healthiness of the food items were similar across all five repetitions. At the individual level, the test-retest reliability for choices in both conditions and healthiness ratings was excellent. However, test-retest reliability for taste ratings was only fair, suggesting that estimates about palatability may vary more from day to day for the same individual.

## Introduction

Food decisions are an integral part of our daily lives. The quality and quantity of the foods we eat have a substantial impact on our health and well-being. There has been a great deal of basic science and clinical research into the determinants and consequences of food choices, as well as numerous efforts to design and validate interventions that lead to healthier eating behaviors^1–5^. Food choice paradigms play a central role in much of this research.

There are several ways food choice paradigms are used in basic and clinical research. Food choice paradigms are a critical tool for investigating the nature of the decision process itself at the behavioral, computational, and neural levels ^6–18^. In addition, they can be used to compare groups ^19–23^, evaluate the effectiveness of a separate intervention (e.g. behavioral, pharmacological, surgical) ^2–4, 16, 24–26^, or even be incorporated into the intervention itself ^27–30^.

Food choice paradigms come in many different flavors. The exact details of the food choice task used in a given study will depend on the hypothesis being investigated and the practical constraints of the experimental setting. For example, in some cases, participants make choices over foods types and quantities in real buffet-like settings, fixed-option ad libitum meals, or snacking opportunities, while in other experiments plastic food replicas, pictures, or text are used to indicate the available options^1^. Often in studies using plastic replicas, pictures, or text, participants are asked to make many different food choices with the understanding that one of those choices will be selected to count, where participants will consume the chosen food “for real”. In general, individuals’ choices over food representations such as pictures are associated with aspects of their actual eating behaviors when measured on the same day or in the future and anthropomorphic measures ^20, 21, 31^, indicating that these paradigms can have a useful degree of ecological and external validity.

Here, we report on decision patterns across five repetitions of a food-picture-based choice paradigm that explicitly cued participants to consider certain food attributes before making their choices on a subset of trials. We refer to this paradigm as the cued-attribute food choice task. Several previous studies have shown that, when participants complete the cued-attribute food choice task a single time, the attribute cues have significant effects on both choice outcomes and brain activity measured by functional magnetic resonance imaging ^7, 13, 32, 33^. Specifically, on trials in which they were cued to consider the healthiness of the food options before making a consumption decision, participants selected healthier items compared to baseline and other cue conditions. However, it is unknown whether repeated experience or practice with the cued-attribute food choice task will lead to changes in choices during health-cued or baseline trials. Does the influence of the health cues increase, decrease, or remain stable with experience and/or time? Does repeated experience with explicitly considering health-related attributes in the cued trials spill over to affect choices in the baseline trials?

We show that the average pattern of choices in the baseline condition as well as the influence of the health cues remain fairly stable over five repetitions of the task across 14 days. Furthermore, the test-retest reliability of individuals’ choices was high across both conditions, consistent with previous reports of high test-retest reliability in other food choice paradigms^34^. Interestingly, the reliability of subjective healthiness ratings was also high, but taste or palatability rating reliability was only fair.

## Results

Figure 1 shows a schematic representation of the food choice task used in this study. Participants were instructed to fast three hours prior the experiment in order to increase the value of foods. In the beginning of the experiment, participants completed a rating phase for 180 images in which they judged in two different phases, how tasty or how healthy they thought each food item to be. In each trial of the subsequent choice task participants have to choose between two food items. Within the choice task, there were two types of decision conditions that differed in the cues that are provided for the participants. In the health-cued condition, subjects are cued to consider the healthiness of the foods while making decisions. In the natural-cued condition, subjects are cued to make decisions naturally based on whatever freely comes to their mind. At the end of each task session one of their choices was randomly selected and participants received and had to eat the chosen food while in the lab. Thus, in both the health-cued and natural-cued conditions, participants had to keep in mind the fact that they may have to eat the food they select at the end of that day’s session.

**Figure 1.**
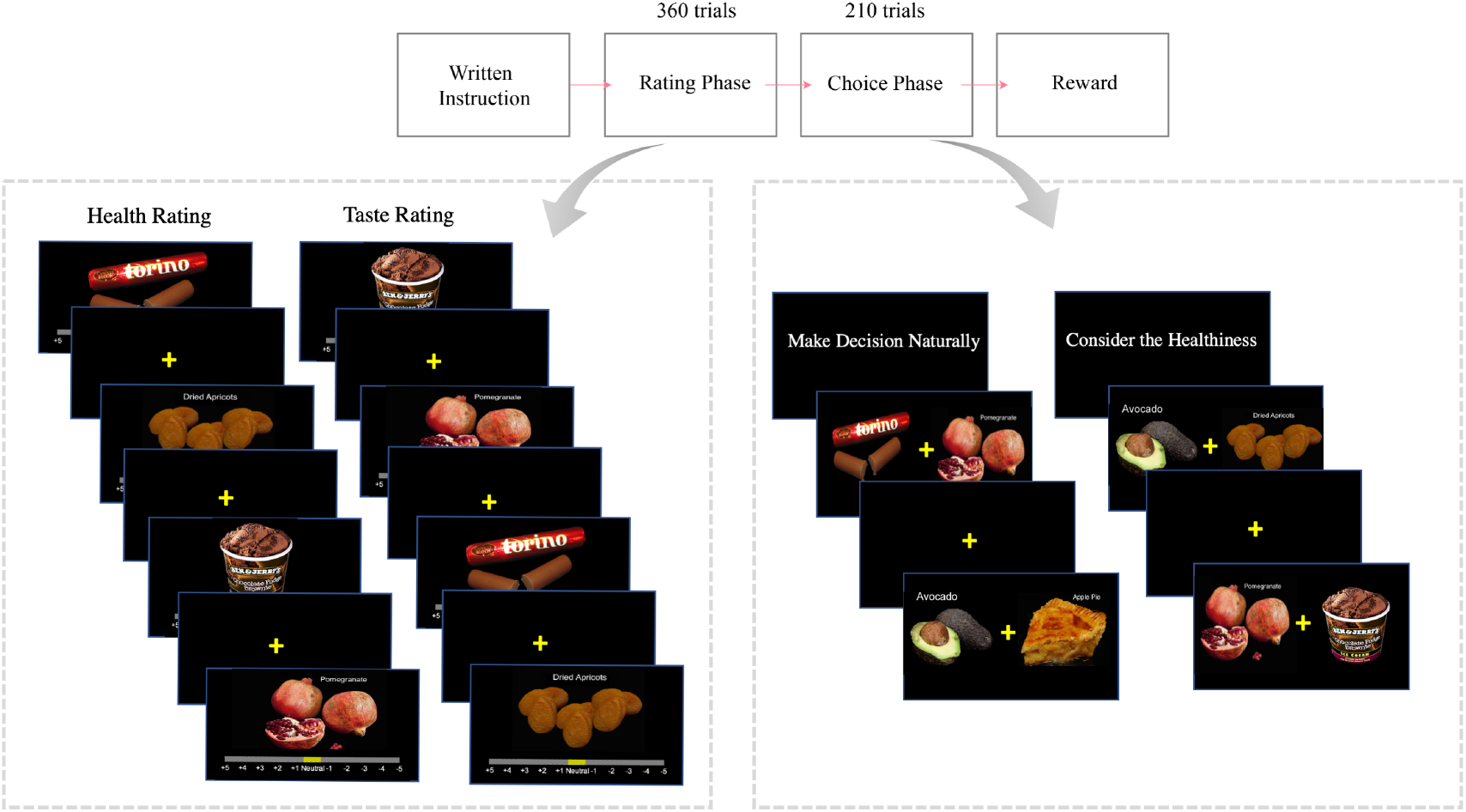
Experiment structure. The experiment had two phases, ratings and choices. In the ratings phase, participants rated the taste and health aspects of the foods using a visual analog scale ranging from −5 to +5. The order of the two ratings was counterbalanced. In the choice task, participants had to choose had to choose one of the two food items to eat at the end of the experiment. Within the choice phase, there were two conditions that differed in the attention cues given to the participants. In the health-cued condition, subjects were cued to consider the healthiness of the foods while making decisions. In the natural-cued condition, subjects were cued to make decisions naturally using whatever features freely came to mind. At the end of the session on each day, one of the participant’s choices was randomly selected and the participant was given the chosen food to eat in the behavioral laboratory.

### Choice outcomes

We examined the effects of attribute (taste, health), context (natural-cued and health-cued condition), and session (1:5) on healthy choice behavior using a Bayesian hierarchical logistic regression. This regression estimated the probability of selecting the healthier of the two food options as a function of the differences in tastiness and healthiness ratings between the foods, the cue condition (natural-cued, health-cued), and the experimental session (1-5). Figure 2 and Table 3 show that, during the baseline (i.e. natural-cued) condition on the first session, the difference in the taste attribute was significantly associated with the choice outcome (1.22, 95% highest density interval (HDI) = [0.94, 1.51]). In contrast, the difference in the healthiness attribute was not significantly related to choice outcomes during natural-cued trials (−0.02, 95% HDI = [−0.20, 0.15]). In session 1, healthier choices were made more often in the health-cued relative to natural-cued condition. There was a significant main effect of health-cued trials (1.52, 95% HDI = [1.04, 1.98]) and significant interactions between the health cues and the differences in both the taste (−0.73, 95% HDI = [−1.02, −0.45]) and healthiness attributes (1.22, 95% HDI = [0.85, 1.60]).

**Figure 2.**
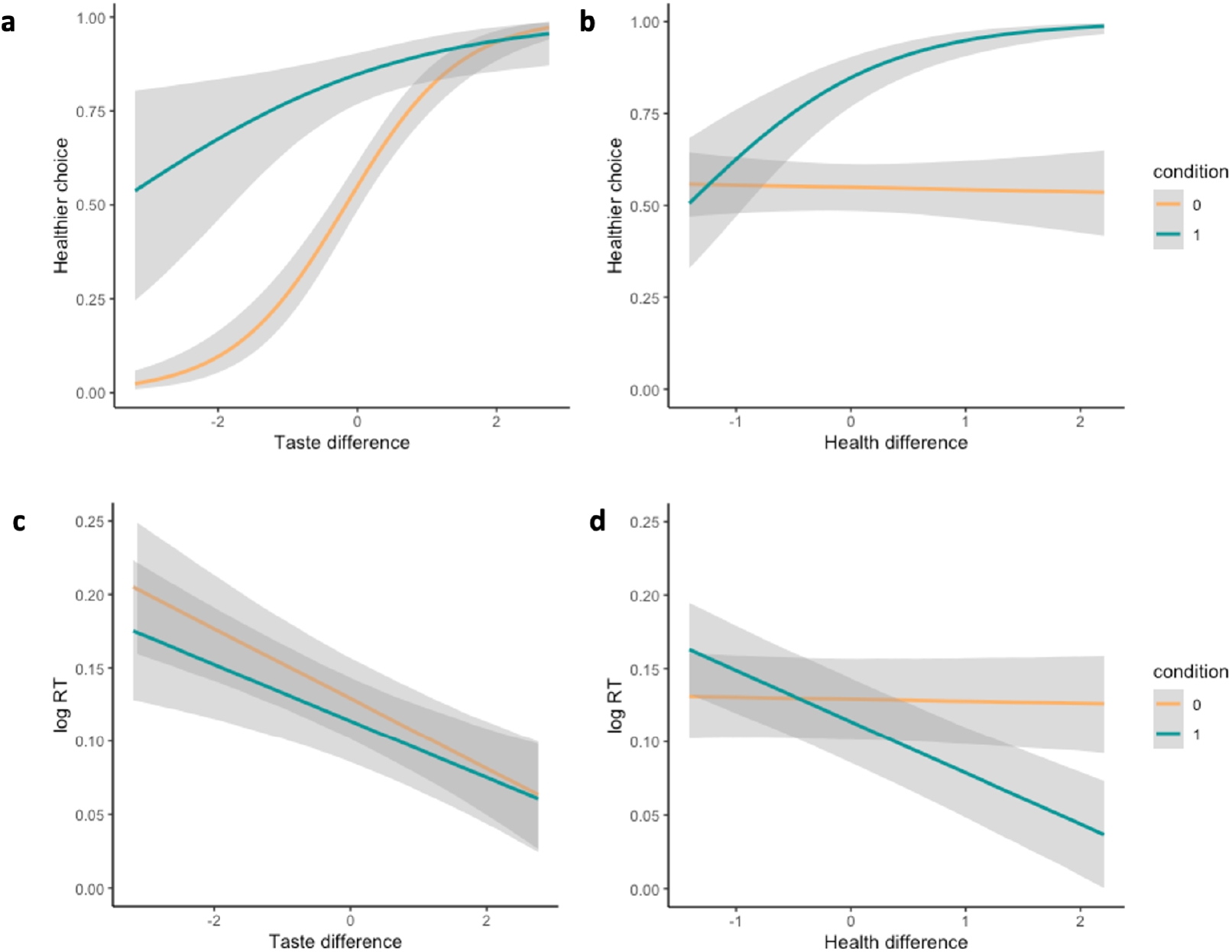
Effect of the taste and healthiness attributes in different conditions on the participants’ choices and response times. **a.** The influence of the taste attributes on whether or not participants made healthier choices was similar in the both natural-(orange lines) and health-cued conditions (green lines), although there is a clear main effect of the health-cue that shifts the curve up toward selecting the healthier option more often. **b.** In contrast, healthiness attributes had almost no influence on choice outcomes in natural-cued trials, but a strong influence in health-cued trials. In both panels **a** and **b**, the y-axis represents the average proportion of healthier choices, while the x-axes show the difference (healthier item minus less healthy item) in the taste or healthiness attributes between the two options, respectively. Panels **c** and **d** show the analogous results replacing the proportion of healthier choices with the logarithm of the response time (logRT) on the y-axis. In all four plots, the grey shaded bars indicate the 95% HDIs for each parameter. Condition 0 is for the natural-cued condition (colored with orange) and condition 1 is for the health-cued condition (colored with green).

**Table 1.** rstDDM Parameters. This table reports the group-level estimates (mean ± standard error (SE)) for each of the rstDDM parameters over 5 sessions in both the natural-cued and health-cued conditions. The parameters are: stochastic component of evidence accumulation (*noise*), starting point bias (*SP bias*; a value of 0.5 indicate no starting point bias), non-decision time (*NDT*), relative start time for health (*RST*; computed as starting time for health minus starting time for taste), weight of taste attribute on drift rate (*taste*), weight of taste attribute on drift rate (*health*), and intercept for drift rate (*Drift bias*). The decision thresholds were held fixed at [−1, 1] when fitting the model.

| *Natural-cued condition* |  |  |  |  |  |
| --- | --- | --- | --- | --- | --- |
| Parameters | session 1 | session 2 | session 3 | session 4 | session 5 |
| Drift bias | 0.02 ± 0.08 | −0.03 ± 0.1 | 0.02 ± 0.11 | 0.08 ± 0.17 | −0.03 ± 0.13 |
| Health weight | −0.13 ± 0.31 | 0.05 ± 0.48 | 0.03 ± 0.47 | −0.01 ± 0.62 | 0 ± 0.49 |
| Taste weight | 0.89 ± 0.48 | 0.69 ± 0.28 | 0.73 ± 0.31 | 0.72 ± 0.39 | 0.87 ± 0.49 |
| Overall drift | 0.78 ± 0.56 | 0.71 ± 0.49 | 0.78 ± 0.35 | 0.79 ± 0.57 | 0.83 ± 0.72 |
| RST | 0.18 ± 0.08 | 0.14 ± 0.13 | 0.02 ± 0.02 | 0.04 ± 0.03 | 0.1 ± 0.06 |
| NDT | 0.6 ± 0.16 | 0.59 ± 0.09 | 0.58 ± 0.15 | 0.54 ± 0.11 | 0.55 ± 0.09 |
| SP bias | 0.49 ± 0.03 | 0.5 ± 0.04 | 0.5 ± 0.03 | 0.48 ± 0.04 | 0.5 ± 0.04 |
| Noise | 1.02 ± 0.13 | 1.02 ± 0.14 | 1.08 ± 0.13 | 1.08 ± 0.16 | 1.09 ± 0.12 |
| *Health-cued condition* |  |  |  |  |  |
| Parameters | session 1 | session 2 | session 3 | session 4 | session 5 |
| Drift bias | 0.05 ± 0.07 | 0.07 ± 0.13 | 0.01 ± 0.12 | 0.08 ± 0.07 | −0.05 ± 0.09 |
| Health weight | 0.9 ± 0.46 | 0.82 ± 0.55 | 0.8 ± 0.5 | 0.68 ± 0.65 | 0.8 ± 0.63 |
| Taste weight | 0.32 ± 0.32 | 0.29 ± 0.23 | 0.3 ± 0.29 | 0.33 ± 0.31 | 0.32 ± 0.44 |
| Overall drift | 1.27 ± 0.51 | 1.19 ± 0.54 | 1.11 ± 0.52 | 1.09 ± 0.63 | 1.07 ± 0.75 |
| RST | −0.03 ± 0.02 | −0.04 ± 0.02 | −0.03 ± 0.03 | −0.02 ± 0.02 | −0.05 ± 0.04 |
| NDT | 0.59 ± 0.13 | 0.57 ± 0.1 | 0.53 ± 0.15 | 0.54 ± 0.1 | 0.52 ± 0.1 |
| SP bias | 0.49 ± 0.03 | 0.48 ± 0.04 | 0.49 ± 0.04 | 0.48 ± 0.04 | 0.51 ± 0.04 |
| Noise | 0.98 ± 0.13 | 1.01 ± 0.12 | 1 ± 0.09 | 1.08 ± 0.12 | 1.03 ± 0.13 |

**Table 2.** Intra-class correlation coefficients (ICC). ICC of rstDDM parameters over 5 sessions for both conditions of natural-cued and health-cued and also for the difference between these two conditions. NDT = non-decision time.

| Parameter | Natural-cued Condition | Health-cued Condition | Health - Natural |
| --- | --- | --- | --- |
| Drift bias | -0.0149 | -0.0679 | -0.129 |
| Health weight | 0.592 | 0.841 | 0.784 |
| Taste weight | 0.53 | 0.696 | 0.541 |
| Overall drift | 0.586 | 0.731 | 0.634 |
| RST | 0.095 | 0.58 | 0.0655 |
| NDT | 0.447 | 0.625 | 0.0277 |
| Bias | 0.0054 | -0.0368 | -0.0264 |
| Noise | 0.522 | 0.477 | -0.00835 |

**Table 3.** Healthier Choice Logistic Regression Model. This table lists all main effects and interactions for the Hierarchical Bayesian logistic regression in Equation 1 (see Methods section). The columns labeled 2.5% and 97.5% indicate the lower and upper bounds of the 95% highest density interval for each parameter, respectively. SD = standard deviation.

| Parameter | Mean | SD | 2.5% | 97.5% |
| --- | --- | --- | --- | --- |
| Intercept | 0.20 | 0.13 | -0.06 | 0.46 |
| td | 1.22 | 0.14 | 0.94 | 1.51 |
| hd | -0.02 | 0.09 | -0.20 | 0.15 |
| condition1 | 1.52 | 0.24 | 1.04 | 1.98 |
| session2 | 0.28 | 0.09 | 0.10 | 0.46 |
| session3 | 0.07 | 0.08 | -0.09 | 0.23 |
| session4 | 0.11 | 0.10 | -0.08 | 0.30 |
| session5 | 0.05 | 0.09 | -0.13 | 0.23 |
| td:condition1 | -0.73 | 0.15 | -1.02 | -0.45 |
| hd:condition1 | 1.22 | 0.19 | 0.85 | 1.60 |
| td:session2 | -0.17 | 0.09 | -0.35 | 0.02 |
| td:session3 | -0.16 | 0.08 | -0.32 | 0.00 |
| td:session4 | -0.21 | 0.09 | -0.40 | -0.03 |
| td:session5 | -0.09 | 0.14 | -0.36 | 0.17 |
| hd:session2 | 0.04 | 0.08 | -0.13 | 0.21 |
| hd:session3 | 0.00 | 0.08 | -0.15 | 0.15 |
| hd:session4 | 0.04 | 0.08 | -0.13 | 0.20 |
| hd:session5 | 0.10 | 0.08 | -0.06 | 0.27 |
| condition1:session2 | -0.39 | 0.13 | -0.65 | -0.12 |
| condition1:session3 | -0.29 | 0.12 | -0.53 | -0.05 |
| condition1:session4 | -0.64 | 0.12 | -0.88 | -0.41 |
| condition1:session5 | -0.25 | 0.13 | -0.49 | 0.00 |
| td:condition1:session2 | 0.16 | 0.11 | -0.06 | 0.38 |
| td:condition1:session3 | 0.14 | 0.11 | -0.07 | 0.35 |
| td:condition1:session4 | 0.20 | 0.11 | -0.01 | 0.41 |
| td:condition1:session5 | 0.20 | 0.12 | -0.03 | 0.42 |
| hd:condition1:session2 | -0.30 | 0.13 | -0.56 | -0.03 |
| hd:condition1:session3 | -0.33 | 0.12 | -0.57 | -0.09 |
| hd:condition1:session4 | -0.54 | 0.12 | -0.77 | -0.31 |
| hd:condition1:session5 | -0.42 | 0.13 | -0.67 | -0.17 |

The overall pattern of behavior observed in session 1, remained similar in sessions 2-5 (Fig 3). There was a slight increase in the average number of healthy choices during the natural condition in session 2 relative to session 1, but no significant difference in natural-cued choices between session 1 and any other session (Table 3). There was a significant increase in the number of healthy choices during the health-cued compared to natural-cued trials within all five sessions, although the difference between health and natural-cued trials decreased slightly after the first session. In addition to choice outcomes, we also tested the repeated subjective ratings for taste and healthiness attributes and found that the group average ratings did not significantly change across sessions (Figure 6).

**Figure 3.**
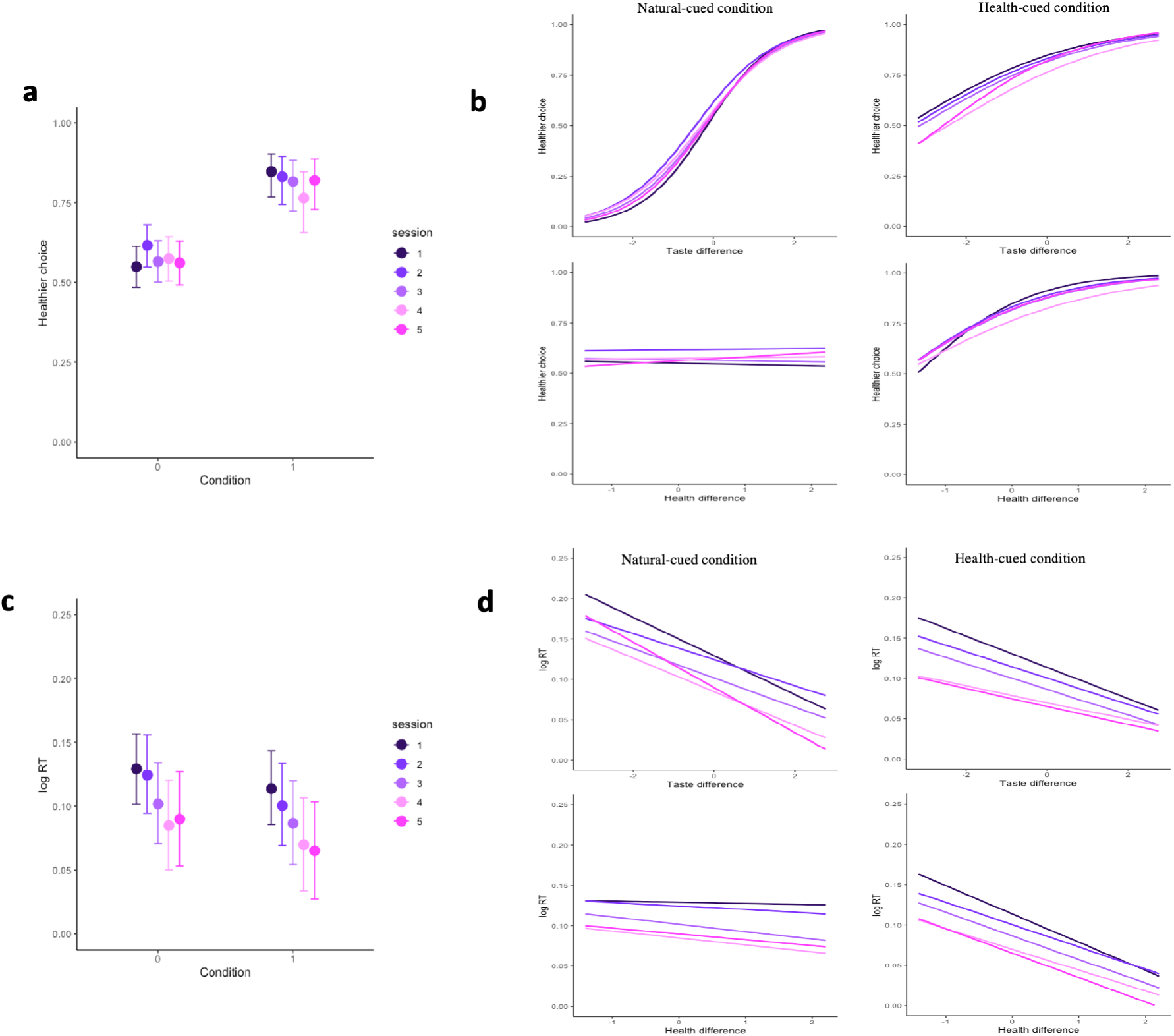
Effects of health-cues and attribute differences on choice outcomes and RTs over the five sessions. Panels **a** and **b** show the proportion of healthier choices on the y-axis as a function of condition (health-cued, natural-cued choice) or attribute differences (computed as healthier item minus less-healthy item) on the x-axis. The results in each session are indicated by the separate colors shown in the legend. **a.** Healthier choices were higher in the health-cued condition compared to the natural-cued condition in all five sessions. **b.** This panel shows how the two attributes (taste and healthiness) relate to choice outcomes in each of the two conditions (natural-cued and health-cued). Panels **c** and **d** are analogous to **a** and **b** except that they show the logarithm of response times on the y-axis as a function of condition or attribute differences on the x-axis. The error bars in both **a** and **c** represent the 95% HDIs.

### Response times

Health cues increased the influence of healthiness attributes on choice response times. In session 1, the healthiness attributes had no significant effect on the response times in the natural condition (0.00, 95% HDI = [−0.01, 0.01]), but did influence response times for health-cued trials (health difference interaction effect = −0.03, 95% HDI = [−0.05, −0.02], Figure 2d). In contrast, taste attributes were significantly associated with response times in both natural and health-cued trials (taste difference main effect = −0.02, 95% HDI = [−0.03,−0.01]). In addition to the interaction between the health-cued condition and health ratings, there was also a main effect condition. Response times were significantly faster overall in the health-cued relative to the natural-cued condition (Figure 2c; main effect of health-cue = −0.02, 95% HDI = [−0.03, 0.00]). The response time patterns in both conditions remained fairly stable across all five sessions, although participants became faster with more experience in the task and there were small changes in the sensitivity to taste and healthiness differences (Figure 2c,d; Tables 4 and 5).

**Table 4.** RT Linear Regression Model, Part I. This table lists all main effects and interactions from the linear regression in Equation 2 (see Methods section) for non-challenge trials. The main effect of challenge trials and its interactions are reported in 5 on the next page for the sake of space. Note that challenge trials are defined as those in which the tastier item is not also the healthier item (i.e. there is a conflict between the taste and health attributes). The regression coefficients in this table represent effects on log(RT) in the subset of trials in which taste and health attributes are aligned. The columns labeled 2.5% and 97.5% indicate the lower and upper bounds of the 95% highest density interval for each parameter, respectively. SD = standard deviation.

| Parameter | Mean | SD | 2.5% | 97.5% |
| --- | --- | --- | --- | --- |
| Intercept | 0.13 | 0.01 | 0.1 | 0.16 |
| td | -0.02 | 0.01 | -0.03 | -0.01 |
| hd | 0 | 0 | -0.01 | 0.01 |
| condition1 | -0.02 | 0.01 | -0.03 | 0 |
| session2 | 0 | 0.01 | -0.02 | 0.01 |
| session3 | -0.03 | 0.01 | -0.05 | -0.01 |
| session4 | -0.04 | 0.01 | -0.07 | -0.02 |
| session5 | -0.04 | 0.01 | -0.07 | -0.01 |
| td:condition1 | 0 | 0.01 | -0.01 | 0.02 |
| hd:condition1 | -0.03 | 0.01 | -0.05 | -0.02 |
| td:session2 | 0.01 | 0.01 | 0 | 0.02 |
| td:session3 | 0.01 | 0.01 | 0 | 0.02 |
| td:session4 | 0 | 0.01 | -0.01 | 0.01 |
| td:session5 | 0 | 0.01 | -0.01 | 0.01 |
| hd:session2 | 0 | 0.01 | -0.01 | 0.01 |
| hd:session3 | -0.01 | 0 | -0.02 | 0 |
| hd:session4 | -0.01 | 0 | -0.02 | 0 |
| hd:session5 | -0.01 | 0.01 | -0.02 | 0 |
| condition1:session2 | -0.01 | 0.01 | -0.03 | 0.01 |
| condition1:session3 | 0 | 0.01 | -0.02 | 0.02 |
| condition1:session4 | 0 | 0.01 | -0.02 | 0.02 |
| condition1:session5 | -0.01 | 0.01 | -0.03 | 0.01 |
| td:condition1:session2 | 0 | 0.01 | -0.02 | 0.01 |
| td:condition1:session3 | 0 | 0.01 | -0.02 | 0.01 |
| td:condition1:session4 | 0.01 | 0.01 | -0.01 | 0.02 |
| td:condition1:session5 | 0.01 | 0.01 | 0 | 0.03 |
| hd:condition1:session2 | 0.01 | 0.01 | 0 | 0.02 |
| hd:condition1:session3 | 0.01 | 0.01 | 0 | 0.03 |
| hd:condition1:session4 | 0.02 | 0.01 | 0 | 0.03 |
| hd:condition1:session5 | 0.01 | 0.01 | 0 | 0.02 |

**Table 5.**
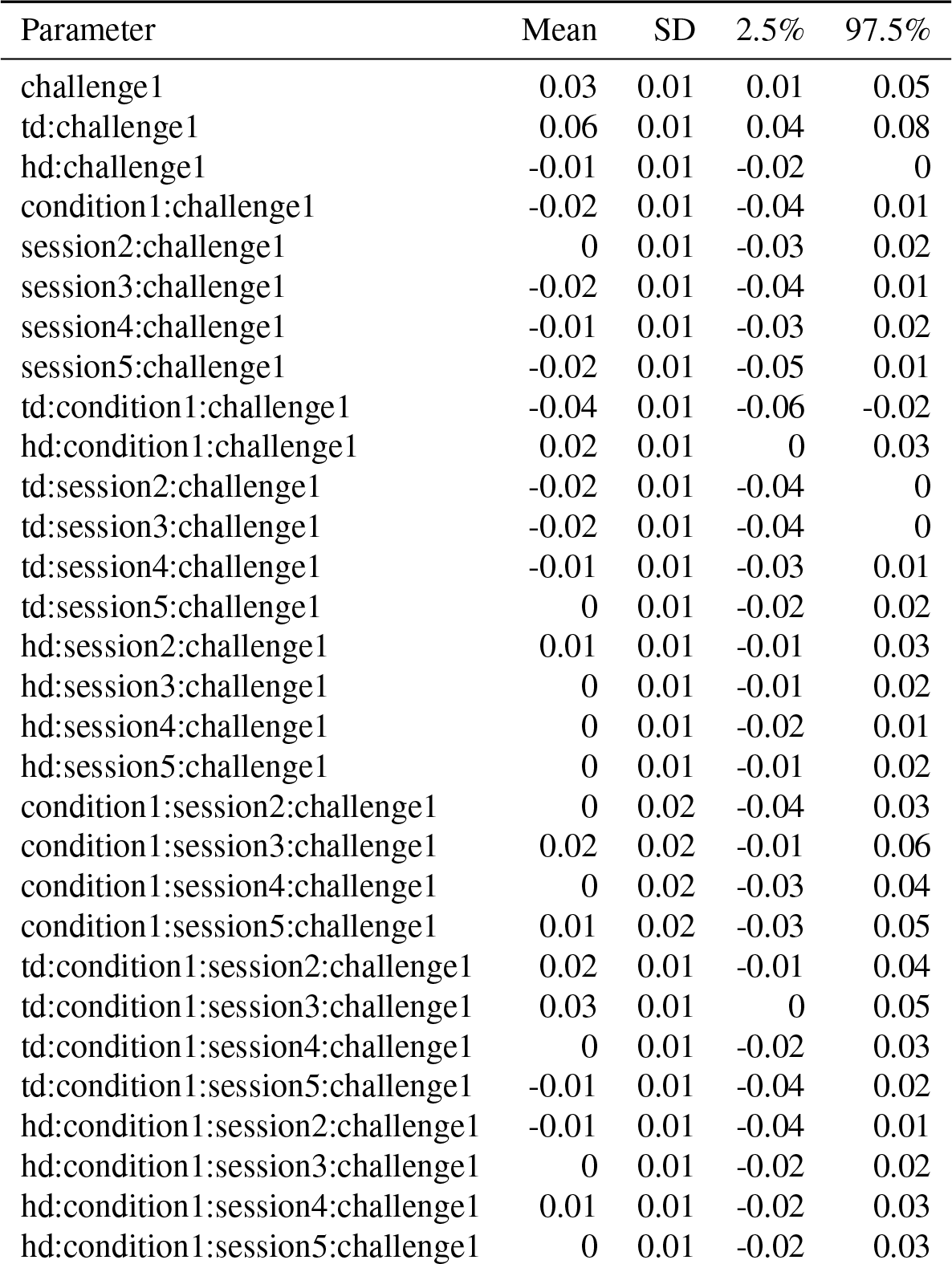
RT Logistic Regression, Part II. This table lists the main effect of challenge trials and its interactions from the linear regression in Equation 2 (see Methods section). Challenge trials are defined as those in which the tastier item is not also the healthier item (i.e. there is a conflict between the taste and health attributes). Thus, the regression coefficients in this table represent effects on log(RT) in the subset of trials in which taste and health attributes are in conflict with regard to which option is preferred. The columns labeled 2.5% and 97.5% indicate the lower and upper bounds of the 95% highest density interval for each parameter, respectively. SD = standard deviation.

| Parameter | Mean | SD | 2.5% | 97.5% |
| --- | --- | --- | --- | --- |
| challenge1 | 0.03 | 0.01 | 0.01 | 0.05 |
| td:challenge1 | 0.06 | 0.01 | 0.04 | 0.08 |
| hd:challenge1 | -0.01 | 0.01 | -0.02 | 0 |
| condition1:challenge1 | -0.02 | 0.01 | -0.04 | 0.01 |
| session2:challenge1 | 0 | 0.01 | -0.03 | 0.02 |
| session3:challenge1 | -0.02 | 0.01 | -0.04 | 0.01 |
| session4:challenge1 | -0.01 | 0.01 | -0.03 | 0.02 |
| session5:challenge1 | -0.02 | 0.01 | -0.05 | 0.01 |
| td:condition1:challenge1 | -0.04 | 0.01 | -0.06 | -0.02 |
| hd:condition1:challenge1 | 0.02 | 0.01 | 0 | 0.03 |
| td:session2:challenge1 | -0.02 | 0.01 | -0.04 | 0 |
| td:session3:challenge1 | -0.02 | 0.01 | -0.04 | 0 |
| td:session4:challenge1 | -0.01 | 0.01 | -0.03 | 0.01 |
| td:session5:challenge1 | 0 | 0.01 | -0.02 | 0.02 |
| hd:session2:challenge1 | 0.01 | 0.01 | -0.01 | 0.03 |
| hd:session3:challenge1 | 0 | 0.01 | -0.01 | 0.02 |
| hd:session4:challenge1 | 0 | 0.01 | -0.02 | 0.01 |
| hd:session5:challenge1 | 0 | 0.01 | -0.01 | 0.02 |
| condition1:session2:challenge1 | 0 | 0.02 | -0.04 | 0.03 |
| condition1:session3:challenge1 | 0.02 | 0.02 | -0.01 | 0.06 |
| condition1:session4:challenge1 | 0 | 0.02 | -0.03 | 0.04 |
| condition1:session5:challenge1 | 0.01 | 0.02 | -0.03 | 0.05 |
| td:condition1:session2:challenge1 | 0.02 | 0.01 | -0.01 | 0.04 |
| td:condition1:session3:challenge1 | 0.03 | 0.01 | 0 | 0.05 |
| td:condition1:session4:challenge1 | 0 | 0.01 | -0.02 | 0.03 |
| td:condition1:session5:challenge1 | -0.01 | 0.01 | -0.04 | 0.02 |
| hd:condition1:session2:challenge1 | -0.01 | 0.01 | -0.04 | 0.01 |
| hd:condition1:session3:challenge1 | 0 | 0.01 | -0.02 | 0.02 |
| hd:condition1:session4:challenge1 | 0.01 | 0.01 | -0.02 | 0.03 |
| hd:condition1:session5:challenge1 | 0 | 0.01 | -0.02 | 0.03 |

### Time-varying diffusion decision model analysis

We used a time-varying diffusion decision model (tDDM) to examine if repeated experience with the cued-attribute food choice task changed the evidence accumulation process assumed to occur during food decisions. The specific tDDM that we used allows for each attribute (here taste and health) to begin to influence the decision process at a different time^?, ?, ?^. We refer to this model as the relative-starting-time DDM (rstDDM). The RST parameter represents the relative advantage in initial processing time for the taste attributes. If RST is positive, then taste-related attributes are considered before healthiness attributes, whereas if RST is negative then healthiness attributes are considered first. The drift weighting parameters in the rstDDM indicate the relative contribution of taste and health to the evidence accumulation process and decision outcome. Figure 4 shows the drift weighting and RST parameters by session and condition (see Table 1 for the full set of parameters).

**Figure 4.**
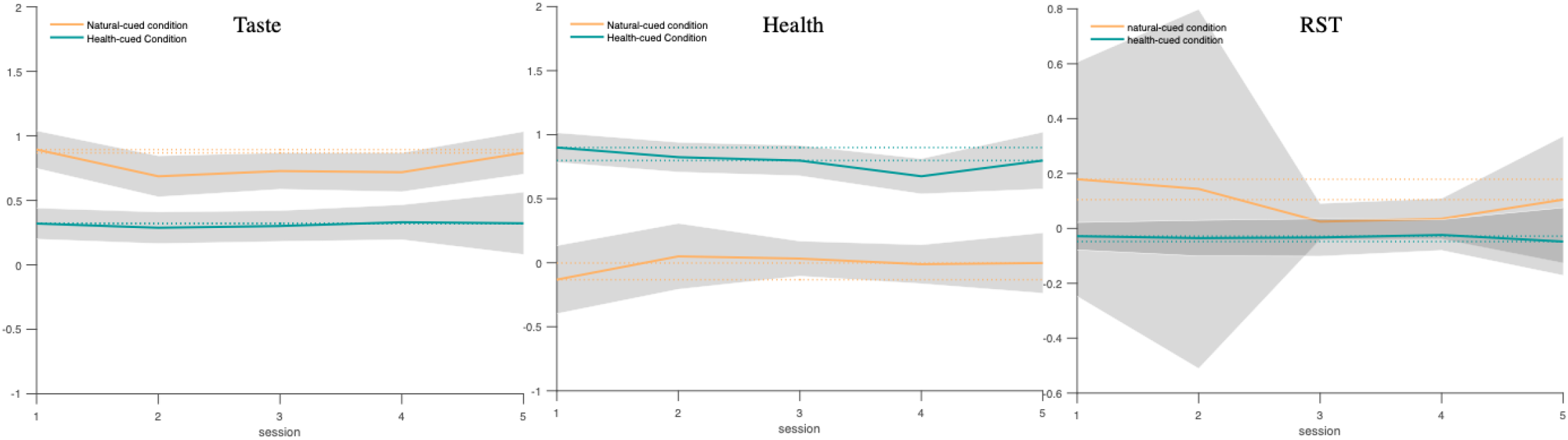
Group-level rstDDM parameters by condition across sessions. These three plots show the overall trends for the taste weight, health weight, and RST parameters in the natural-cued and health-cued conditions across sessions. The taste weight was higher in the natural-cued condition compared to the health-cued condition in all sessions (left panel). The health weight was higher in the health-cued condition compared to the natural-cued condition in all sessions (middle panel). The RST parameter was qualitatively lower (i.e. healthiness was considered earlier) in the health-cued condition compared to the natural-cued condition in all sessions (right panel), although there was substantial individual variability in the RST parameter during the natural-cued trials. All group-level parameters other than the RST parameter for natural-cued trials were quite stable and did not differ with repeated experience across sessions. The shaded bars indicate the 95% HDIs for each parameter. The natural-cued condition is indicated by orange color and health-cued condition is indicated by green color.

Health cues had a significant and stable influence on rstDDM parameters across all five sessions. On average, participants weighted taste attributes more and considered them sooner than health attributes in the natural-cued condition during the first-session (taste weight: 0.8940, 95% HDI = [0.8200, 0.9660], health weight: −0.132, 95% HDI = [−0.275, −0.009], rst: 0.1790, 95% HDI = [0.021, 0.448]), and this pattern held across all sessions (Fig. 4). However, during the health-cued trials health weights were higher than in the natural trials (0.8120, 95% HDI = [0.7520, 0.8780]), while in contrast taste weights showed the opposite pattern and were significantly higher in the natural-cued condition relative to the health-cued condition (0.4670, 95% HDI = [0.4180, 0.5150]). Comparing the health and taste weights within each condition revealed that the health weight was significantly higher than taste weight in the health-cued condition (0.4870, HDI= [0.4290, 0.5430]), and taste weight were significantly higher than health weight in the natural-cued condition (0.7910, HDI= [0.7240, 0.8620]). The total contribution of taste plus health attributes to the drift rate was significantly higher in the health-cued than natural-cued condition (0.3450, 95% HDI = [0.2760, 0.4130]), which is consistent with the pattern of faster mean RTs in the health-cued trials shown in Figure 3. In addition to weights, health cues also significantly changed the relative-starting times for each attribute within the decision process. In the first session, the RST was also smaller (i.e. healthiness was considered earlier) in the health-cued condition compared to the natural-cued condition (−0.1310, 95% HDI = [−0.2680, −0.0530]). Health-cues continued to promote the consideration of health attributes before taste across all five sessions. Across sessions, the consideration start times for healthiness and tastiness in natural cued trials became more similar (i.e. RST was closer to 0), and less variable across individuals (Fig. 4).

In general individual participants’ rstDDM parameters were fairly stable across sessions. Figure 5 shows that the ranges (*max – min*) for the parameter estimates across the five sessions were fairly small, with the notable exception of the RST for natural-cued trials. Comparisons across conditions showed that the ranges were significantly smaller in the health-cued condition than in the natural-cued condition (Taste: *tstat* = −3.1051, *pvalue* = 0.0052, Health: *tstat* = −2.4076, *pvalue* = 0.0249, RST: *tstat* = −14.5627, *pvalue* = 8.8958*e* – 13).

**Figure 5.**
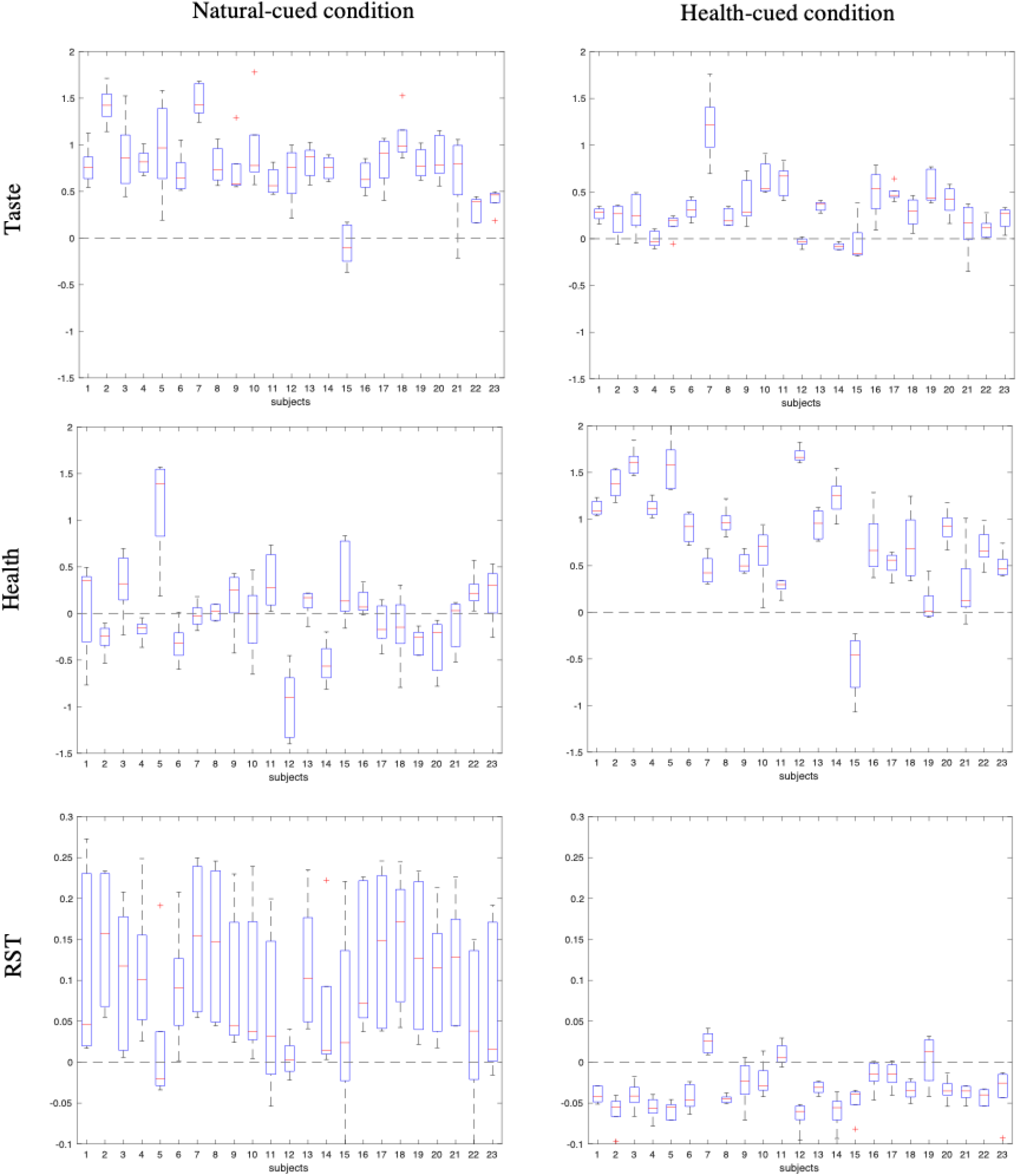
rstDDM parameters stability. The boxplots show that the values of most subject-level parameters didn’t change much across the 5 sessions. However, the changes in attribute weights on the drift rate over time/experience were significantly greater in the natural-cued condition compared to the health-cued condition. We address the high variability of the *RST* parameters in the natural-cued condition further in the discussion section.

**Figure 6.**
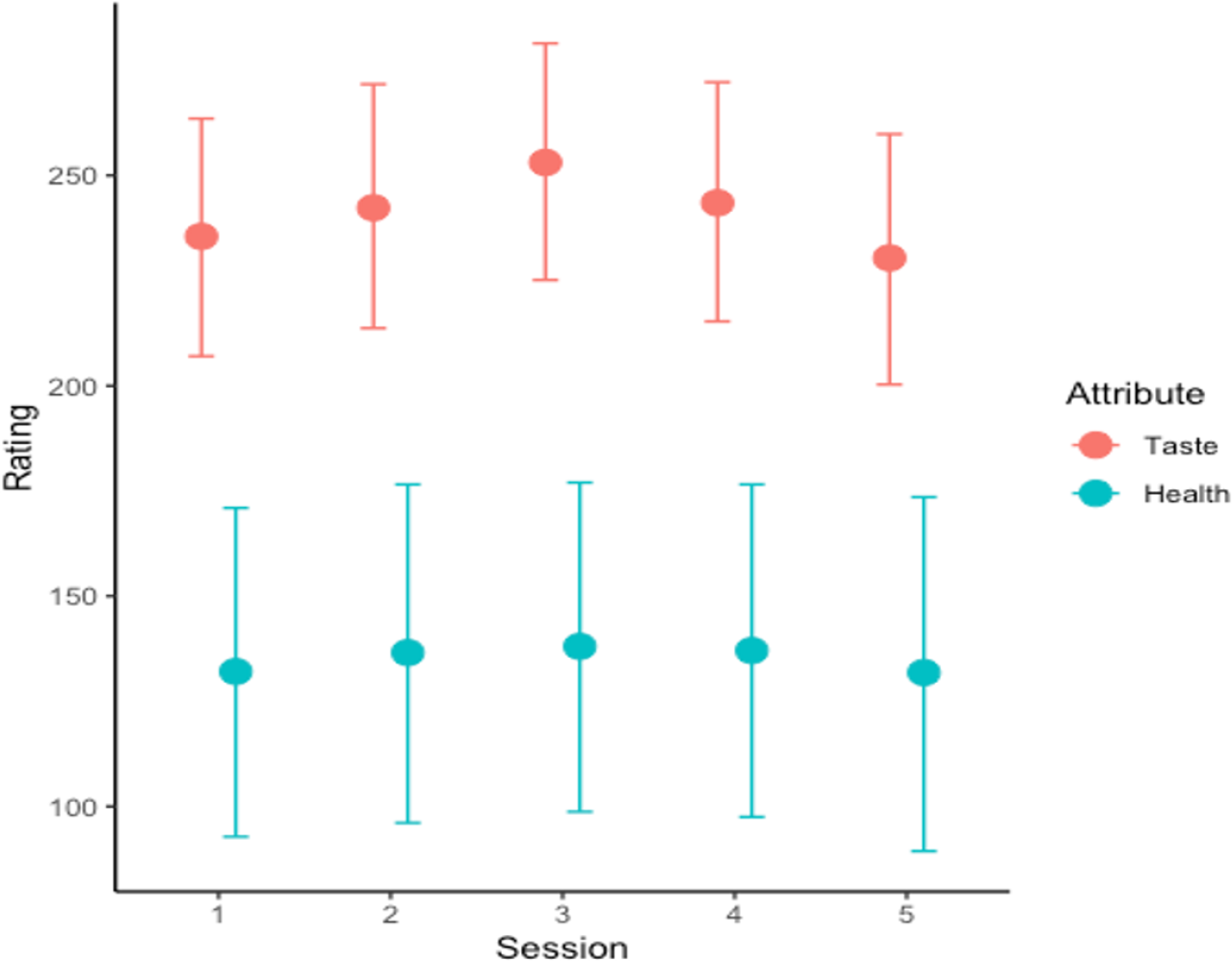
Ratings over sessions. The ratings didn’t significantly changed over sessions. Dimension 1 is for taste weight and is indicated by red color, and dimension 2 is for health weight and is indicated by light green. The error bars indicate standard error of means.

### Test-retest reliability at the individual level

Having found that group average decisions within the cued-attribute food choice task are fairly stable over 14 days and 5 repetitions of the task, we next tested the reliability at the individual level. Specifically, we computed the intra-class correlation coefficients (ICC), for three different task measures: 1) frequency of healthier choices, 2) subjective ratings of taste and healthiness, and 3) rstDDM parameters (health/taste weight, and RST). In the text below we interpret the ICC values as: Poor (< 0.40), Fair (0.4 – 0.6), Good (0.6 – 0.75), and Excellent (0.75 – 1.0)^?^.

Individuals’ choices showed excellent test-retest reliability across the five sessions. The frequency of healthier choices showed excellent reliability across the 5 sessions both within the natural- and health-cued conditions separately and as a difference score between conditions. (natural-cued condition, *ICC*(*A*, 1) = 0.808, *F*(22, 86.8) = 23.3, *p* = 1.74*e* – 27, 95%-Confidence Interval: 0.688 < *ICC* < 0.901, health-cued condition, *ICC*(*A*, 1) = 0.812, *F*(22, 91.9) = 22.6, *p* = 6.06*e* – 28,95%-Confidence Interval: 0.695 < *ICC* < 0.903, difference between conditions, *ICC*(*A*, 1) = 0.812, *F*(22, 89) = 23.5, *p* = 4.79*e* – 28, 95%-Confidence Interval: 0.694 < *ICC* < 0.902).

The test-retest reliability for participants’ subjective healthiness and tastiness ratings was excellent and fair, respectively. Note that participants did not face the exact same choices in each session, but rather similar types of choices drawn from the same set of 180 food items. In addition to making choices, the participants rated each of the 180 food items for taste and healthiness in sessions 1 and 5. They rated a subset of 60 food items on taste and health in all five sessions. We computed the ICC both for full set of 180 and subset of 60 items. Rating reliability at the individual level was excellent for health and fair for taste ratings (60 items in all 5 sessions, health: 0.808±0.171, taste: 0.579±0.184, Table 6, 180 items in sessions 1 and 5, health: 0.778±0.216, taste: 0.532±0.190, Table 6). We also analysed rating reliability as a function of food image rather than individual. At the food image level, we found that reliability was fair for both health and taste ratings (across five sessions: health: 0.552±0.146, taste: 0.574±0.120, Table 7, between session 1 and 5 alone: health: 0.468±0.251, taste: 0.516±0.197, Table 8).

**Table 6.**
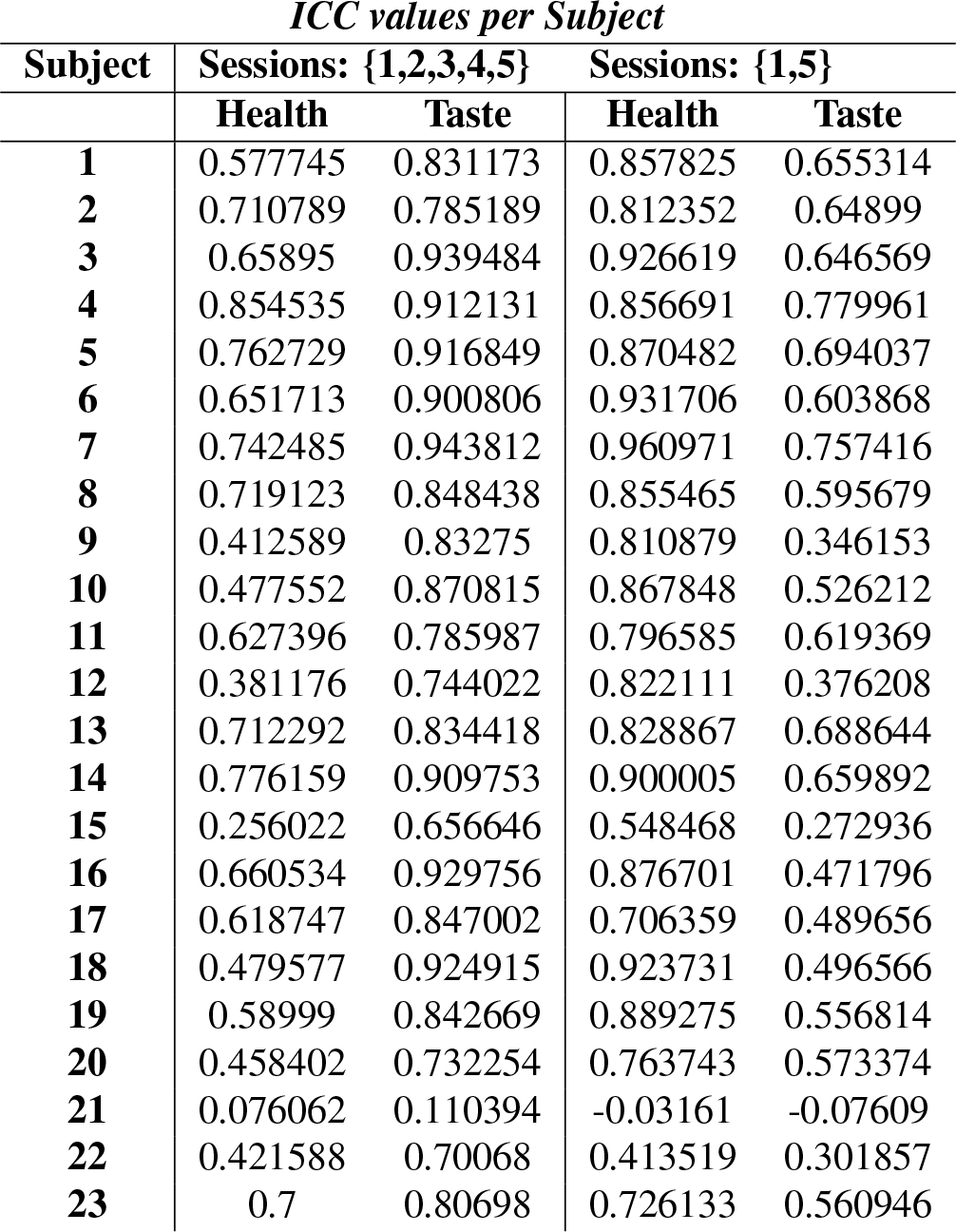
ICC values per subject for the subset of 60 food images rated in all 5 sessions/days (left) and the full set of 180 food images rated in sessions 1 and 5 (right).

| *ICC values per Subject* |  |  |  |  |
| --- | --- | --- | --- | --- |
| **Subject** | **Sessions: {1,2,3,4,5}** |  | **Sessions: {1,5}** |  |
|  | **Health** | **Taste** | **Health** | **Taste** |
| **1** | 0.577745 | 0.831173 | 0.857825 | 0.655314 |
| **2** | 0.710789 | 0.785189 | 0.812352 | 0.64899 |
| **3** | 0.65895 | 0.939484 | 0.926619 | 0.646569 |
| **4** | 0.854535 | 0.912131 | 0.856691 | 0.779961 |
| **5** | 0.762729 | 0.916849 | 0.870482 | 0.694037 |
| **6** | 0.651713 | 0.900806 | 0.931706 | 0.603868 |
| **7** | 0.742485 | 0.943812 | 0.960971 | 0.757416 |
| **8** | 0.719123 | 0.848438 | 0.855465 | 0.595679 |
| **9** | 0.412589 | 0.83275 | 0.810879 | 0.346153 |
| **10** | 0.477552 | 0.870815 | 0.867848 | 0.526212 |
| **11** | 0.627396 | 0.785987 | 0.796585 | 0.619369 |
| **12** | 0.381176 | 0.744022 | 0.822111 | 0.376208 |
| **13** | 0.712292 | 0.834418 | 0.828867 | 0.688644 |
| **14** | 0.776159 | 0.909753 | 0.900005 | 0.659892 |
| **15** | 0.256022 | 0.656646 | 0.548468 | 0.272936 |
| **16** | 0.660534 | 0.929756 | 0.876701 | 0.471796 |
| **17** | 0.618747 | 0.847002 | 0.706359 | 0.489656 |
| **18** | 0.479577 | 0.924915 | 0.923731 | 0.496566 |
| **19** | 0.58999 | 0.842669 | 0.889275 | 0.556814 |
| **20** | 0.458402 | 0.732254 | 0.763743 | 0.573374 |
| **21** | 0.076062 | 0.110394 | -0.03161 | -0.07609 |
| **22** | 0.421588 | 0.70068 | 0.413519 | 0.301857 |
| **23** | 0.7 | 0.80698 | 0.726133 | 0.560946 |

**Table 7.** ICC values per image across for the subset of 60 items rated in all five sessions.

| *ICC values per food image across 5 sessions* |  |  |  |  |  |
| --- | --- | --- | --- | --- | --- |
| **Images** | **Health** | **Taste** | **Images** | **Health** | **Taste** |
| **1** | 0.548981 | 0.360017 | **31** | 0.415038 | 0.374519 |
| **2** | 0.839613 | 0.678043 | **32** | 0.601435 | 0.510712 |
| **3** | 0.530135 | 0.435967 | **33** | 0.381346 | 0.500656 |
| **4** | 0.499733 | 0.524707 | **34** | 0.567766 | 0.599284 |
| **5** | 0.53434 | 0.570703 | **35** | 0.715889 | 0.812522 |
| **6** | 0.450081 | 0.623053 | **36** | 0.646967 | 0.600088 |
| **7** | 0.718801 | 0.671339 | **37** | 0.739166 | 0.641482 |
| **8** | 0.453639 | 0.597161 | **38** | 0.65807 | 0.70777 |
| **9** | 0.882878 | 0.594293 | **39** | 0.569736 | 0.664011 |
| **10** | 0.662751 | 0.51526 | **40** | 0.667879 | 0.656011 |
| **11** | 0.494073 | 0.709408 | **41** | 0.375358 | 0.564663 |
| **12** | 0.580517 | 0.716292 | **42** | 0.535164 | 0.432079 |
| **13** | 0.341531 | 0.412314 | **43** | 0.741156 | 0.538456 |
| **14** | 0.533415 | 0.618469 | **44** | 0.403696 | 0.384647 |
| **15** | 0.502964 | 0.394111 | **45** | 0.568851 | 0.431005 |
| **16** | 0.387244 | 0.533008 | **46** | 0.569612 | 0.561693 |
| **17** | 0.204656 | 0.397756 | **47** | 0.437377 | 0.585203 |
| **18** | 0.330214 | 0.433892 | **48** | 0.619319 | 0.648813 |
| **19** | 0.786993 | 0.66636 | **49** | 0.492178 | 0.58252 |
| **20** | 0.63245 | 0.681147 | **50** | 0.255104 | 0.533144 |
| **21** | 0.435499 | 0.568086 | **51** | 0.372384 | 0.506155 |
| **22** | 0.73386 | 0.656252 | **52** | 0.520931 | 0.599399 |
| **23** | 0.544993 | 0.486982 | **53** | 0.60342 | 0.570972 |
| **24** | 0.611825 | 0.669386 | **54** | 0.631005 | 0.699958 |
| **25** | 0.64508 | 0.658395 | **55** | 0.695773 | 0.711901 |
| **26** | 0.739376 | 0.596953 | **56** | 0.707249 | 0.678658 |
| **27** | 0.473611 | 0.770296 | **57** | 0.644811 | 0.740512 |
| **28** | 0.690062 | 0.491442 | **58** | 0.626182 | 0.719768 |
| **29** | 0.424566 | 0.710365 | **59** | 0.309981 | 0.519247 |
| **30** | 0.29652 | 0.170655 | **60** | 0.574235 | 0.492516 |

**Table 8.**
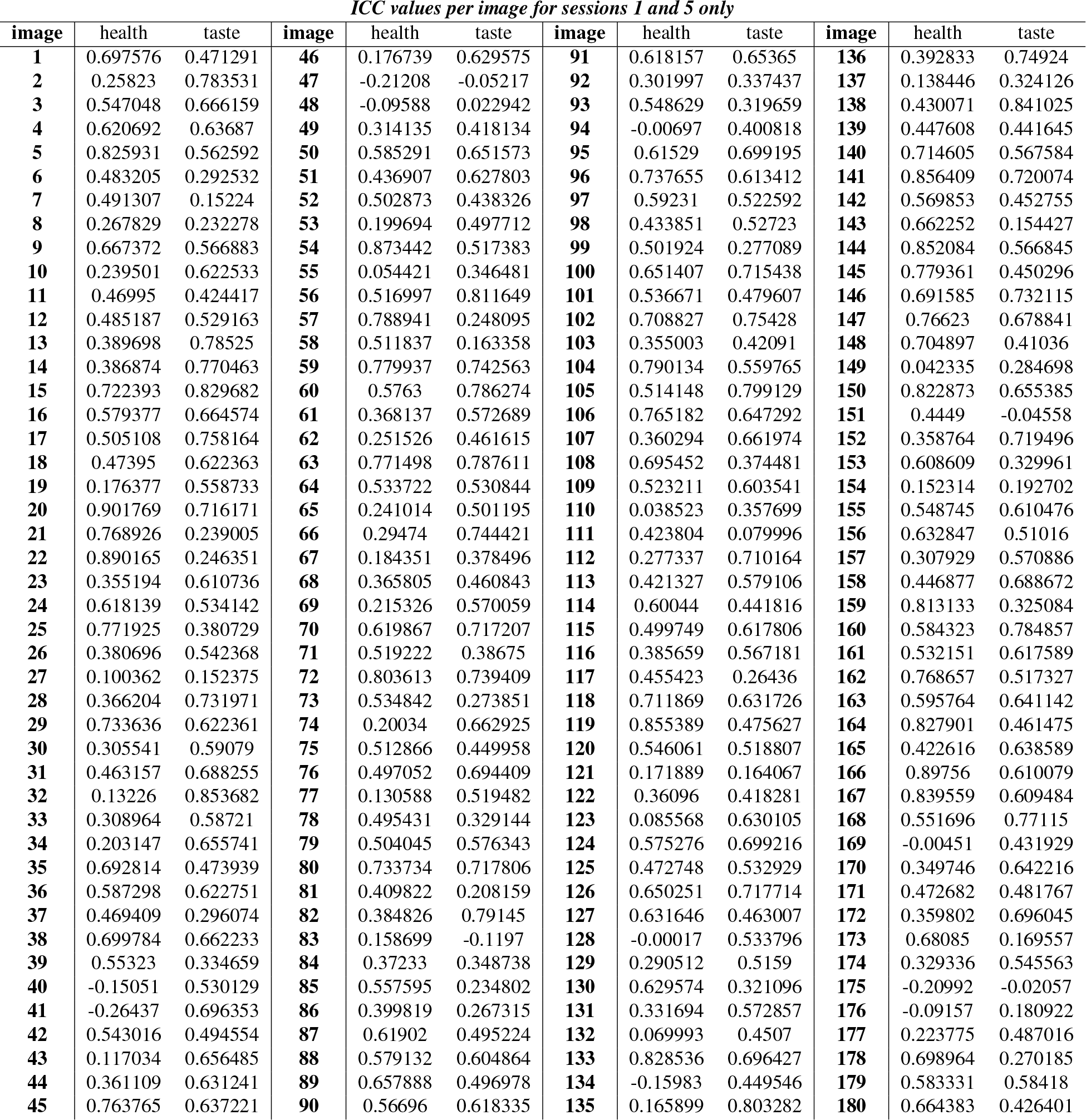
ICC values per image across sessions 1 and 5 for the full set of 180 images.

Lastly, we computed the test-retest reliability of the rstDDM parameters fit to the participants’ choice outcomes and response times. Table 2 reports the ICC values for each parameter by condition. The reliability of the parameters ranged from excellent to good in the health-cued condition. However, the reliability of the same parameters fit to the same individuals was only fair to poor in the natural-cued condition. As a consequence of the lower reliability of the parameters in the natural-cued condition, the reliability of the difference scores for taste and health weights is also lower.

## Discussion

Previous research has shown that food choice tasks using photographic images as stimuli have good test-retest reliability when participants make choices naturally (i.e. without explicit instructions about how to choose or what attributes to consider) ^34^. However, unlike the previously tested paradigms, there are clear differences in the ways the same participant will make choices across the different conditions within the cued-attribute food choice task ^7, 13, 32, 33^. We tested whether these intra-individual differences changed over the course of 14 days and five repetitions of the cued-attribute food choice task. Overall, choice outcome patterns were stable and had excellent test-retest reliability within and across the decision conditions over the five task repetitions. Interestingly, healthiness ratings had excellent reliability, but the reliability for taste ratings was only fair. This suggests that a individuals’ estimates of the tastiness of food items may vary more from day to day than their opinions about healthiness.

Although choice outcomes had excellent reliability in both the natural and health-cued task conditions, the reliability of rstDDM parameters differed across conditions. A much stronger reliance on tastiness than healthiness attributes may have led to the lower test-retest reliability for rstDDM parameters fit to the natural-cued choices. Both the rstDDM and the logistic regression analyses showed that, on average, healthiness had little influence on food choice outcomes during natural-cued choices in our sample of participants. Figures 3b and 4 show that, holding tastiness fixed, the influence of healthiness on food decision is approximately zero at the group level. This is not too surprising given that the individuals were recruited because they consumed sweet or savory snacks on a regular basis, but attempting to eat a healthy diet in daily life was not an inclusion criterion. Thus, concerns about the palatability of the food items dominated the decision process in natural trials. The near-zero influence of healthiness on food choices in natural trials means that the rstDDM has little power to accurately identify the relative attribute weighting and consideration start timing parameters (*RST*).

The rstDDM model we fit to the data can accurately identify and distinguish between attribute weights and *RST* parameters when both weights are non-zero ^?, ?^, but (near) zero weights are a problem for it. Either a sufficiently low drift weighting coefficient or long delay in consideration starting time can effectively eliminate the influence of an attribute on choice outcomes within the rstDDM framework. The excellent reliability of the drift weighing parameters and good reliability of the *RST* parameter in the health-cued trials proves that the rstDDM can yield highly reliable parameter estimates if fit to a suitable set of data. Thus, the lower reliability of the rstDDM in the natural-cued trials serves to demonstrate the importance of checking that the data conform to the expectations and requirements of a given modeling approach. If one of the drift weights from the rstDDM is close to zero for a given data set, then one is probably better off fitting the more common version of the DDM ^?^, which fixes the *RST* parameter to zero and only estimates the drift weights (see Table 9 for the full set of parameters, and Table 10 for the reliability values).

**Table 9.**
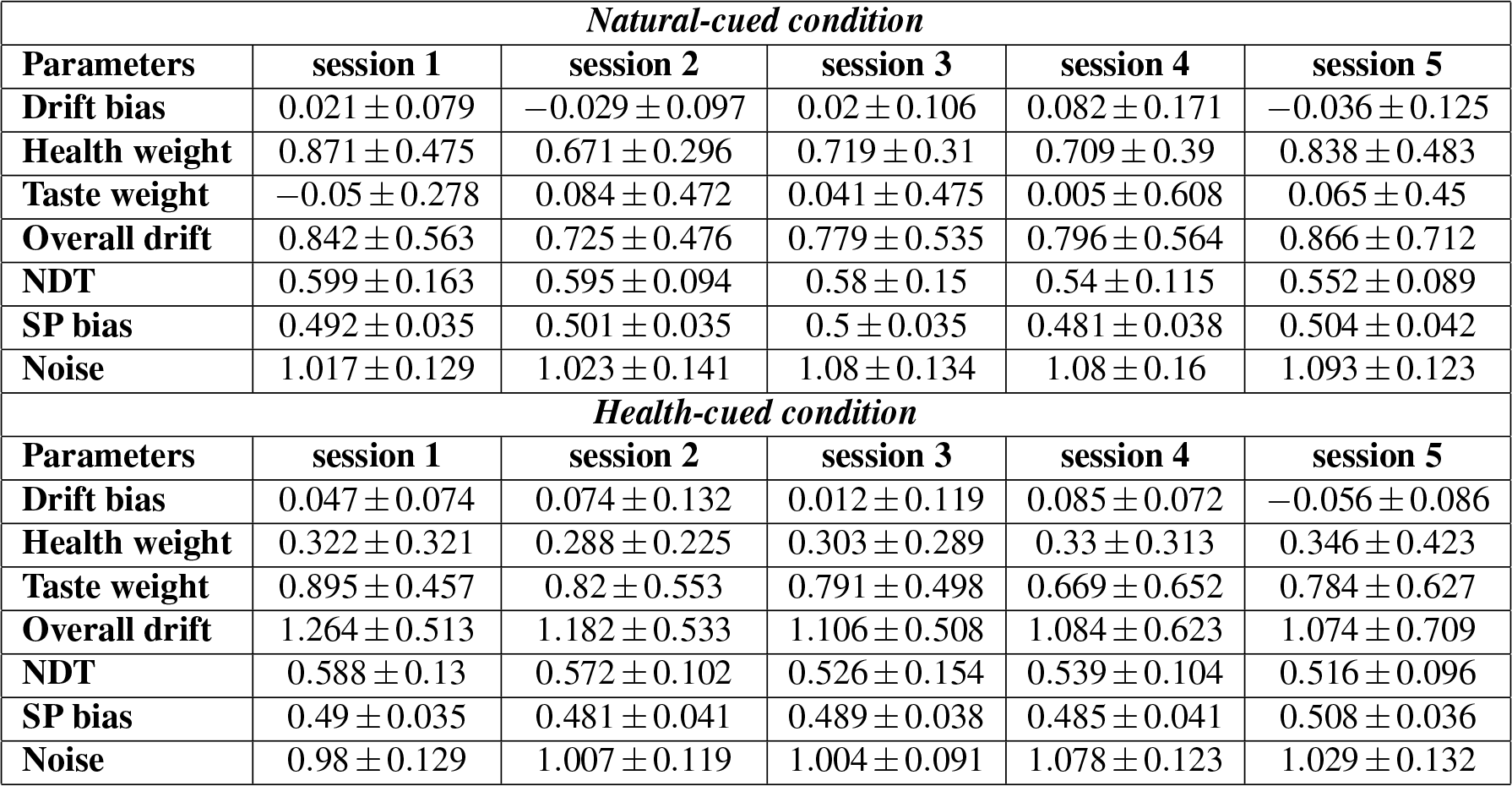
Standard-DDM Parameters. This table reports the group-level estimates (mean ± standard error (SE)) for each of the standard-DDM parameters over 5 sessions in both the natural-cued and health-cued conditions. The parameters are: stochastic component of evidence accumulation (*noise*), starting point bias (*SP bias*; a value of 0.5 indicate no starting point bias), non-decision time (*NDT*), weight of taste attribute on drift rate (*taste*), weight of taste attribute on drift rate (*health*), and intercept for drift rate (*Drift bias*). The decision thresholds were held fixed at [−1, 1] when fitting the model.

| *Natural-cued condition* |  |  |  |  |  |
| --- | --- | --- | --- | --- | --- |
| Parameters | session 1 | session 2 | session 3 | session 4 | session 5 |
| Drift bias | 0.021 ± 0.079 | −0.029 ± 0.097 | 0.02 ± 0.106 | 0.082 ± 0.171 | −0.036 ± 0.125 |
| Health weight | 0.871 ± 0.475 | 0.671 ± 0.296 | 0.719 ± 0.31 | 0.709 ± 0.39 | 0.838 ± 0.483 |
| Taste weight | −0.05 ± 0.278 | 0.084 ± 0.472 | 0.041 ± 0.475 | 0.005 ± 0.608 | 0.065 ± 0.45 |
| Overall drift | 0.842 ± 0.563 | 0.725 ± 0.476 | 0.779 ± 0.535 | 0.796 ± 0.564 | 0.866 ± 0.712 |
| NDT | 0.599 ± 0.163 | 0.595 ± 0.094 | 0.58 ± 0.15 | 0.54 ± 0.115 | 0.552 ± 0.089 |
| SP bias | 0.492 ± 0.035 | 0.501 ± 0.035 | 0.5 ± 0.035 | 0.481 ± 0.038 | 0.504 ± 0.042 |
| Noise | 1.017 ± 0.129 | 1.023 ± 0.141 | 1.08 ± 0.134 | 1.08 ± 0.16 | 1.093 ± 0.123 |
| *Health-cued condition* |  |  |  |  |  |
| Parameters | session 1 | session 2 | session 3 | session 4 | session 5 |
| Drift bias | 0.047 ± 0.074 | 0.074 ± 0.132 | 0.012 ± 0.119 | 0.085 ± 0.072 | −0.056 ± 0.086 |
| Health weight | 0.322 ± 0.321 | 0.288 ± 0.225 | 0.303 ± 0.289 | 0.33 ± 0.313 | 0.346 ± 0.423 |
| Taste weight | 0.895 ± 0.457 | 0.82 ± 0.553 | 0.791 ± 0.498 | 0.669 ± 0.652 | 0.784 ± 0.627 |
| Overall drift | 1.264 ± 0.513 | 1.182 ± 0.533 | 1.106 ± 0.508 | 1.084 ± 0.623 | 1.074 ± 0.709 |
| NDT | 0.588 ± 0.13 | 0.572 ± 0.102 | 0.526 ± 0.154 | 0.539 ± 0.104 | 0.516 ± 0.096 |
| SP bias | 0.49 ± 0.035 | 0.481 ± 0.041 | 0.489 ± 0.038 | 0.485 ± 0.041 | 0.508 ± 0.036 |
| Noise | 0.98 ± 0.129 | 1.007 ± 0.119 | 1.004 ± 0.091 | 1.078 ± 0.123 | 1.029 ± 0.132 |

**Table 10.** Intra-class correlation coefficients (ICC). ICC of Standard-DDM parameters over 5 sessions for both conditions of natural-cued and health-cued and also for the difference between these two conditions. NDT = non-decision time.

| Parameter | Natural-cued Condition | Health-cued Condition | Health - Natural |
| --- | --- | --- | --- |
| Drift bias | -0.0146 | -0.067 | -0.13 |
| Health weight | 0.538 | 0.718 | 0.541 |
| Taste weight | 0.671 | 0.838 | 0.808 |
| Overall drift | 0.615 | 0.742 | 0.648 |
| NDT | 0.446 | 0.625 | 0.0278 |
| SP bias | 0.00853 | -0.037 | -0.0248 |
| Noise | 0.519 | 0.479 | -0.0114 |

Given its high level of test-retest reliability, the cued-attribute food choice task appears to be well suited to serve as an outcome measure for cognitive, physical, or pharmacological interventions that target food consumption decision processes. However, it is important to note that here we have only tested healthy young adults. Further tests in individuals with obesity, eating disorders, and other conditions are warranted.

The cued-attribute food choice task explicitly prompts participants to incorporate health-related attributes into their decision process. In combination with functional magnetic resonance imaging, the health-cued condition within this task has been shown to engage prefrontal brain activity patterns similar to those observed during endogenous dietary self-control in individuals who successful lost or maintained their desire weight ^7, 19, 32, 33^. Thus, the cued attribute food choice task may be a means of testing how well an intervention shifts externally cued health-promoting decision processes into an individual’s natural, default behavior.

## Methods

### Participants

Twenty-three participants completed the five sessions of the experiment across 14 days. All the participants (10 female, ages between 18 – 30 years) gave written informed consent for their participation in accordance with the regulations of the Zurich Cantonal Ethics commission. They received a flat fee in addition to their food reward for the time they spent for the task. We screened participants by email and phone to ensure that they did not follow any specific food diet, and they consumed sweet and/or savory snacks regularly. We also rechecked the exclusion criteria on each day of testing. All participants were healthy and without any current or recent psychiatric, metabolic or neurological illness. On the day of the study, participants sent us a photograph of the meal they consumed 3 hours before their appointment as an indication that they followed our instructions to eat a small meal and then fast for 3 hours before coming into the lab. All five laboratory visits took place between 17:15 and 19:15. The first visit occurred on a Wednesday, visits 2:3 took place on the Monday, Wednesday, and Friday of the following week. The fifth and final visit took place on a Wednesday two weeks after the initial visit. All procedures were approved by and performed in accordance with the guidelines and regulations of the Zurich Cantonal Ethics commission.

### Behavioral Task

Participants were asked to eat a small meal three hours before their appointment and consume nothing but water in the meantime, in order to be hungry and thereby increase the value of the foods during the experiment. The task involved two sequential phases of food rating and food choice, which participants performed in all 5 sessions. Before each phase, subjects performed a short training session to become familiar with the task. In the food rating phase, participants judged how tasty or healthy each food item is in two different blocks. The ratings were done on continuous scale with anchors of −5 and +5 at each end. The order of taste and health rating blocks were counterbalanced, and the order of images within each block was randomized. In the first and last sessions (1 and 5), the ratings were done over all 180 food images, but in the middle sessions (2-4), the ratings were done for a subset of 60 food images whose average ratings in previous studies spanned the total ranges for taste and healthiness. We reduced the size of the rating set in order to save time in sessions 2:4 and maximize participant retention.

In the food choice phase, subjects had to choose which of two food items they would eat at the end of the experiment. The food pairs were constructed so that choosing the healthier item often required forgoing the subjectively tastier one, and we refer to these cases as challenge trials. One of the participant’s decisions were randomly selected and implemented after the task. There was a 30-minute waiting period following the end of the session during which participants ate their selected food item. This 30-minute waiting period made the decisions more meaningful because participants did not have access to other food sources at that time, and they had not eaten for approximately 4.5 hours at that point.

Within the choice phase, there were two types of conditions that differed in the attention cues that are provided for the participants. In the health-cue condition, subjects were cued to consider the healthiness of the foods while making decisions. In this condition they were supposed to choose the healthier of the foods as often as they could, while keeping in mind that they would potentially have to eat the food they choose. In the natural-cue condition, subjects were cued to make decisions naturally based on whatever freely came to their minds. The food choice task consisted of 3 runs, with each run consisting of 5 or 6 blocks, 210 trials in total. The order of condition blocks was pseudo-randomized across subjects. Each subject faced 9 health-cue condition blocks (110.6±5.2 trials) and 8 natural-cue condition blocks (99.3±5.2 trials).

Choices were presented on the screen for 3 s and there was a jittered inter-trial interval was of 2–6 s. After the task was completed, one trial randomly was selected to be realized for each participant according to his or her choices. The participants stayed in the laboratory for 30 min to eat their food reward.

### Computational tools

#### Logistic regression for choice outcome

We ran a Bayesian hierarchical logistic regression analysis with the *brms* package^?, ?^ in *R*^?^, to estimate the participants choice. We used the uninformative priors default in the brms package. We estimated the probability of choosing the healthier food option, as a function of the difference in taste ratings (td) and health ratings (hd) between the healthier and less-healthy options, condition (natural-, and health-cued conditions), and session (5 sessions). The model formula in *brms* syntax was:

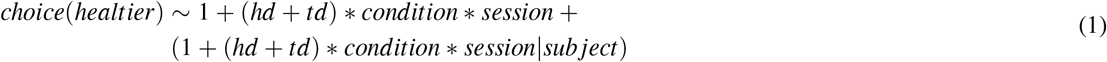

Note that asterisks indicate interaction terms and thus the model included all main effects and interactions between regressors except for interactions between taste and health differences. The first line lists the regressors and interactions that were estimated at the group or population level, and the final portion on the second lined inside parentheses shows those that were estimated at the subject level. In this case, all terms were estimated at both levels.

#### Linear regression for response time

We ran another Bayesian hierarchical regression analysis with the *brms* package in *R*, to estimate the response times (logarithm) as a function of difference in taste ratings and health ratings between two competing options, condition (natural-, and health-cued conditions), session (5 sessions), challenge type (when there is a conflict between taste and health and they have similar ratings). The health and taste ratings were z-scored, and all other variables were categorical. We used the uninformative priors default in the brms package. The model in *brms* syntax was:

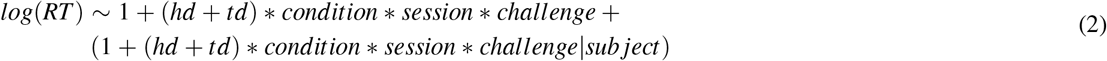

Note that asterisks indicate interaction terms and thus the model included all main effects and interactions between regressors except for interactions between taste and health differences. The first line lists the regressors and interactions that were estimated at the group or population level, and the final portion on the second lined inside parentheses shows those that were estimated at the subject level. In this case, all terms were estimated at both levels.

#### Sequential sampling model

We used the relative-starting-time DDM (rstDDM) model to estimate the response times and choice outcomes of the participants. This model was previously tested and validated for binary food choices ^?^. In this work, we used the hierarchical Bayesian fitting procedure described here: https://github.com/galombardi/method_HtSSM_aDDM. We fit each session and condition separately. The following seven free parameters were estimated in each fitting: stochastic component of evidence accumulation (*noise*), starting point bias (*SP bias*), non-decision time (*NDT*), relative start time for health (*RST*), weight of taste attribute on drift rate (*ω_taste_*), weight of health attribute on drift rate (*ω_health_*), and intercept for drift rate (*Drift bias*). The decision thresholds were fixed to [−1, 1].

#### Test-retest Reliability(ICC)

To determine the test-retest reliability of the task across the five sessions, we computed intra-class correlation coefficients (ICC). We computed ICCs for three different task dimensions including frequency of healthier choices, and DDM parameters (focusing on drift bias, health/taste weight, and RST), and participants’ ratings (health and taste). All of the variables were continuous, therefore we used the *icc* function in the *irr* library in R^?^ with the following configuration: *icc*(*data*, *model* = “*twoway*”, type = “*agreement*”, *unit* = “*single*”, *r*0 = 0, *conf.level* = 0.95). We follow Shrout and Fleiss^?^ and interpret the ICC values according to the following convention: Poor (< 0.40), Fair (0.4 – 0.6), Good (0.6 – 0.75), and Excellent (0.75 – 1.0).

## Acknowledgements

This work was supported by a grant from the Swiss National Science Foundation (SNSF; grant number 32003B_166566).

## Author contributions statement

ARB and TAH designed the study. ARB and ZB acquired the data. ZB analyzed the data with input from ARB and TAH. TAH and ZB wrote the paper with input from ARB.

## Data availability

The datasets analyzed for this study can be found in the https://osf.io/bhjpz/.

## Additional information

### Ratings

The group mean values for subjective ratings of taste and healthiness did not significantly change across sessions (Figure 6).

